# 3D printed bioreactor enabling the physiological culture and the study of pathological processes in large vessels

**DOI:** 10.1101/2022.02.01.477871

**Authors:** Rolando S Matos, Davide Maselli, John H McVey, Christian Heiss, Paola Campagnolo

## Abstract

Routine cardiovascular interventions such as balloon angioplasty, result in vascular activation and remodeling, often requiring re-intervention. *2D in vitro* culture models and small animal experiments have enabled the discovery of important molecular and cellular pathways involved in this process, however the clinical translation of these results is often underwhelming. There is a critical need for an *ex vivo* model representative of the human vascular physiology and encompassing the complexity of the vascular wall and the physical forces regulating its function. Vascular bioreactors for *ex vivo* culture of large vessels are viable alternatives, but their custom-made design and insufficient characterization often hinders the reproducibility of the experiments.

The objective of the study was to design and validate a novel 3D printed cost-efficient and versatile perfusion system, capable of sustaining the viability and functionality of large porcine arteries for 7 days and enabling monitoring of post-injury remodeling.

MultiJet Fusion 3D printing technology was used to engineer the ‘*EasyFlow*’ insert, converting a conventional 50 ml centrifuge tube into a mini bioreactor. Porcine carotid arteries either left untreated or injured with a conventional angioplasty balloon, were cultured under pulsatile flow for up to 7 days. Pressure, heart rate, medium viscosity and shear conditions were adjusted to represent the typical arterial physiology. Tissue viability, cell activation and matrix remodeling were analyzed by immunohistochemistry, and vascular function was monitored by duplex ultrasound.

Physiological blood flow conditions in the *EasyFlow* bioreactor preserved endothelial coverage and smooth muscle organization and extracellular matrix structure in the vessel wall, as compared to static culture. Injured arteries presented hallmarks of early remodeling, such as intimal denudation, smooth muscle cell disarray and media/adventitia activation in flow culture. Duplex ultrasound confirmed physiological hemodynamic conditions, dose-dependent vasodilator response to nitroglycerin in untreated vessels and impaired dilator response in angioplastied vessels. We here validate a low-cost, robust and reproducible system to study vascular physiopathology, laying the basis for future investigations into the pathological remodeling of blood vessels and creating a platform to test novel therapies and devices *ex vivo*, in a patient relevant system.

## Introduction

Vascular disease is often caused by narrowing or occlusion of blood vessels leading to decreased blood supply to important organs like the heart or brain, or to the extremities. Revascularization procedures are common interventions aimed at re-establishing blood supply to compromised tissues. Typically, a guidewire is passed through an accessible artery to reach the narrowed area and a balloon is deployed to reopen the lumen. While the immediate effect on blood flow is readily achieved, the intervention damages the blood vessel wall which triggers a cascade of acute inflammatory and regenerative responses that result in intimal hyperplasia and vascular remodeling (1). The vascular remodeling commonly leads to re-stenosis and ultimately occlusion of the intervened vessels (2). In order to maintain patency in the long term, many times additional devices like stents or balloons coated with anti-proliferative drugs are used on the intervened vessels. Clearly, a better understanding of the processes that underly the vascular response to injury and remodeling is key to develop effective treatments to improve long-term patency of revascularization procedures (1).

Largely, experiments aimed at investigating these mechanisms are conducted in small animals (mice, rats, rabbits), upon mechanical injury of a large artery or interposition of a stent (3). Besides the obvious ethical concerns, small rodent experiments have limited predictive capacity, resulting in the development of sub-optimal therapeutic strategies, leaving this critical medical need unmet (4). *In vitro* strategies for the study of vascular remodeling focus on 2D culture of vascular cells (typically smooth muscle cells) or the static culture of rings obtained from blood vessels, the latter importantly incorporates an element of intact extracellular matrix component, which is critical in the remodeling process (for an example (5)). Interestingly, some groups have demonstrated the successful use of flow bioreactors simulating the *in vivo* hemodynamic conditions, to culture blood vessels *ex vivo* and study vascular pathophysiology (6–9). This pioneering work has demonstrated the importance and feasibility of a functional and physiological *ex vivo* approach to vascular remodeling; however, these studies present some limitations that reduce their translational power and reproducibility, which we aim to address with the present work.

In this study, we developed an open-source 3D printed and economical bioreactor (EasyFlow) which enables the multiplex culture of blood vessels from large animals in small volumes of medium. Furthermore, we have optimized conditions of flow, shear stress, pressure, pulsatility and viscosity to closely mimic the physiological forces applying to an artery *in vivo*. We comprehensively characterized the effect of physiological culture and endovascular injury on the survival, activation and function of vascular resident cells by histology, immunofluorescence and by Doppler ultrasound vasoreactivity measurements.

This work will demonstrate the reproducible application of *ex vivo* bioreactor culture for large animal blood vessels, enabling the long-term culture of healthy vessels and the study of pathological mechanisms. Importantly, the open-source nature of our design enables effortless transfer of the system to other labs and reproducibility, promoting the reduction of the use of animals in vascular studies.

## Materials and Methods

### Ethics and sample preparation

Carotid arteries were obtained from 4-6 weeks old piglet at The Pirbright Institute, Pirbright, UK. Animal procedures were carried out under the Home Office Animals (Scientific Procedures) Act (1986) and approved by the Animal Welfare and Ethical Review Board (AWERB) of The Pirbright Institute. The animals were housed in accordance with the Code of Practice for the Housing and Care of Animals Bred.

Pigs were euthanized by an overdose of 10ml pentobarbital (Dolethal 200mg/ml solution for injection, Vetoquinol UK Ltd). All procedures were conducted by Personal License holders who were trained and competent and under the Project License PPL70/8852.

Upon exsanguination the neck was opened from the chest cavity to the base of the skull to expose the common carotid artery separating into the right and left carotid artery. Arteries were excised using a no-touch technique to minimize stress to the vascular tissue. Fresh tissue was immediately placed in pre cooled transport media Dulbecco’s Modified Eagle’s Medium (DMEM) + 20% fetal bovine serum (FBS) + 2% penicillin and streptomycin (P/S) + 1% Amphotericin B (Amp). Tissue was washed two times in transport media and placed on ice for transportation.

Following arrival tissue was transferred into laminar hood where further preparation took place. The excess connective tissue was removed in a Petri dish using precision surgical equipment, avoiding any strain to the vessel. Once cleaned, the vessel was placed in new container and washed in transport media (2x 20 min at 4°C), followed by short term storage in DMEM + 10% FBS + 1% P/S + 1% Amp at 4°C.

For each preparation, a sample was collected at the time of preparation, as a control.

### EasyFlow system set up

Details of the EasyFlow bioreactor system are presented in **Figure 1** and in the Result section in detail.

**Figure 1.**
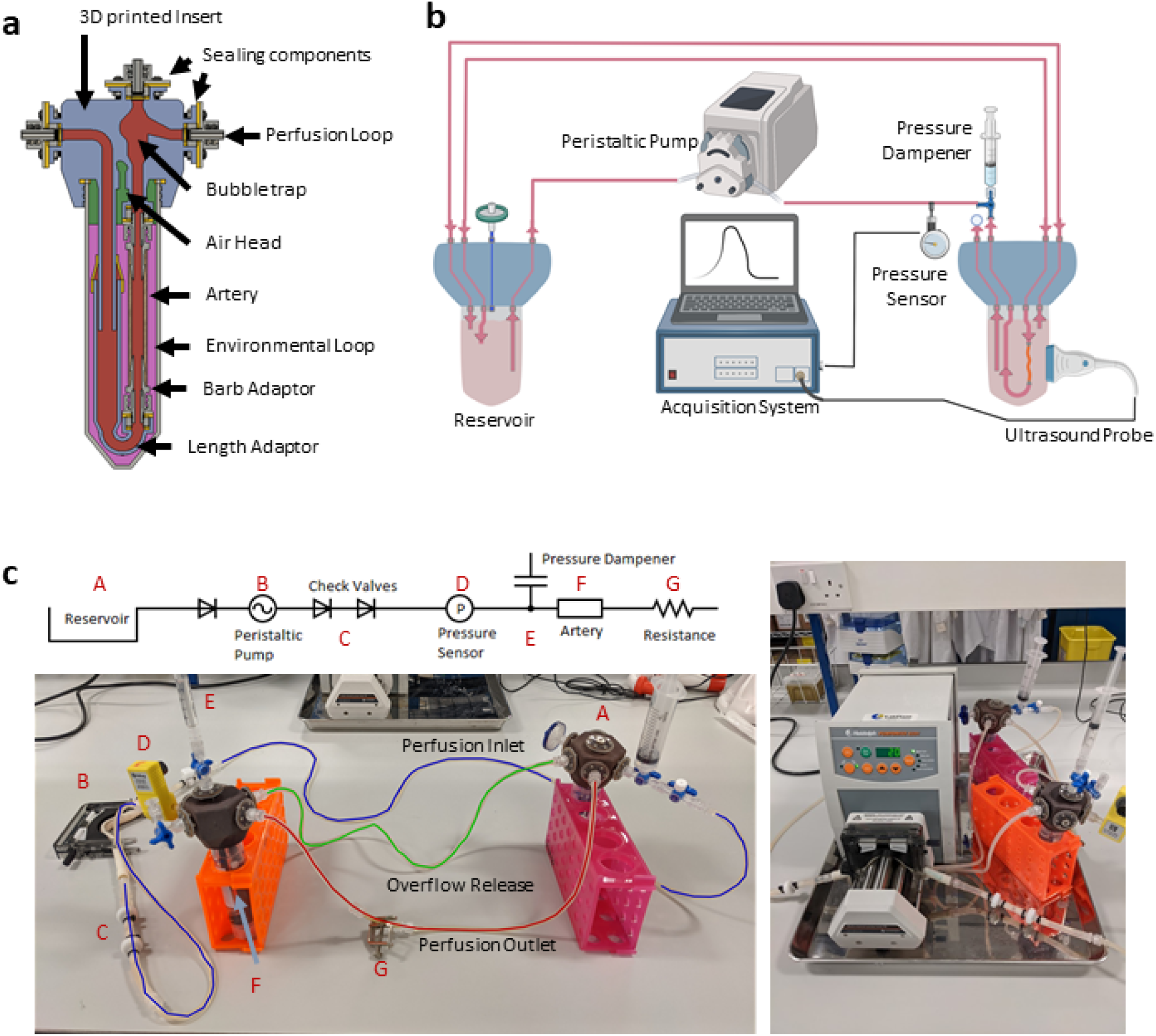
EasyFlow system characteristics and setup. Cross-sectional view of the EasyFlow insert visualizing the inner structure and interconnectivity of the reaction space contained within a 50 mL centrifuge tube (a). Schematic of the perfusion loop which includes the EasyFlow bioreactor accommodating the porcine carotid artery, a reservoir, the peristaltic pump, and acquisition system (b). A simplified connection map and true image of system identifying the key elements of the perfusion loop (c).

The EasyFlow insert consists of a 3D printed part that fits as a lid of a 50 ml standard centrifuge tube. The EasyFlow insert presents 6 inlet and outlet ports, creating two separate circulations inside and outside the blood vessel, which is lodged in its adaptor (**Figure 1a**). The insert was 3D printed by MultiJet fusion of Polyamide 12 (HP 3D High Reusability PA 12) according to the 3D design generated in Fusion 360 (Autodesk^®^ Fusion 360™ 2.0.5818).

The bioreactor system is composed by three main parts: a peristaltic pump (Heidolph, PD 5201, 523-52010-00-2) mounting a 4-cassette head (Heidolph, Multichannel Pump Head C8, 524-40810-00), an EasyFlow insert lodging the blood vessel sample and an equivalent EasyFlow functioning as media reservoir (**Figure 1b**). The two inserts are connected through a series of tubes. Most of the components are connected with transparent silicon tubing (RS Components, RS PRO Silicone Transparent Silicone Tubing, 3.2mm Bore Size, 667-8444). Larger bore peristaltic pump compatible tubing (RS Component, Verderflex Verderprene Yellow Process Tubing, 6.4mm Bore Size, 125-4042) were used in the pump head. Unidirectional flow is ensured by check valves (Cole-Parmer, Masterflex PVDF and Viton® Barbed Check Valve, 3/16-inch, WZ-98766-50) added downstream the peristaltic pump, and pressure is stabilized by a pressure dampener system created with a syringe loaded with liquid and air. To increase baseline pressure low bore tubing (RS Components, RS PRO Silicone Transparent Silicone Tubing, 0.8mm Bore Size, 667-8432) was used downstream of the artery. To regulate resistance, an additional clamp (Cole-Parmer, Flow Control Clamps, 08028-00) was applied downstream of the resistance tubing. Sterile gas exchange is enabled by the presence of outlets mounted with sterility filters (Sarstedt, Syringe filter, Filtropur S, PES, pore size: 0.2 μm, 83.1826.001) (**Figure 1c**).

The system was monitored by pressure sensors downstream of the sample-containing EasyFlow (Parker, SciLog SciPres Pressure Sensor Luer Connection, SCIPRE080699PSX), and through repeated Duplex ultrasound imaging performed with Logiq V2 (GE Healthcare).

To assure the aseptic environment necessary for the long-term incubation of tissues, all components were sterilized by autoclaving. Elements of the circulation system were assembled under a laminar flow hood and primed with *perfusion media* (DMEM + 10% FBS + 1% P/S + 1 % Amp + 3% Dextran). The circulation system was inspected for leaks, and any bubble trapped in the system was removed by flowing media across the system.

Previously harvested and cleaned tissue was then placed in the EasyFlow unit. Depending on the internal diameter of the vessel segment, an adequate barb connector was selected to facilitate the connection to the perfusion system, typically for porcine carotid arteries 2.4 or 1.6mm ID connectors were used (Cole-Parmer, Female luer x 5/32” hose barb adapter, 45501-06). The length adaptor within EasyFlow is ultimately adjusted to create a minimal stretch on the longitudinal axis (**Figure 1a**).

Once the tissue was secured using surgical vessel bond (Vessel Loop Maxi, SKU: AR-722), and residual air was removed from the newly assembled segment before sealing the system and transferring it to the incubator (5% CO_2_ at 37°C). Circulation was connected to the peristaltic pump (**Figure 1c**) and primed overnight with low flow conditions (5 rpm). This allows system equilibrium and troubleshooting while minimizing unnecessary strain on the tissue. Flow was gradually increased (+1 rpm every 60 min) until the setpoint is achieved (14 rpm).

### Balloon injury procedure

A standard peripheral artery angioplasty balloon catheter (Cook Mediacal, Advance® 35LP Low 4 mm) was inserted into the artery. With a standard indeflator pump the balloon was inflated with phosphate buffer saline (PBS) to 6 atm and the pressure was maintained for 3 min. After deflation, the balloon was extracted, the tissue rinsed and placed in the perfusion system for culture, or static culture as a control.

### Artery culture in flow or static conditions

Healthy and balloon injured arteries were cultured for 7 days in static or flow conditions. In static culture, 2-4 mm long tissue segments are submerged in medium (DMEM + 10% FBS + 1% P/S + 1% Amp) in Petri dishes, and incubated at 5% CO_2_, 37°C. Every three days 50% of the media was replaced with fresh media.

Perfusion cultures were performed using the EasyFlow bioreactor system, as described above, on segments of 2-4 cm of length. The system is filled with ~100 ml of *perfusion media* and the peristaltic pump is set at the final speed of 14 rpm (equivalent to 56 pulse/min). In our system set up, this rate corresponds to a 42 ml/min volumetric flow, resulting in a calculated flow velocity of ~40 cm/s within the artery. To assure that the tissue is not subject to extreme pressure conditions generated by the peristaltic movement, a pressure dampener was introduced to maintain fluctuations within the physiological range (60-120 mmHg).

Samples in flow were monitored at day 3 and 7 by Duplex ultrasound imaging. At the end of the culture period, all samples were processed for histology and *en face* staining.

### Doppler ultrasound imaging and vasoreactivity assay

Ultrasound images of the incubated arteries were obtained with a standard clinical vascular ultrasound machine with a 10 MHz linear array transducer (LOGIQ V2, GE Healthcare). Briefly, ultrasound gel was applied to the outside of the bioreactor and longitudinal images obtained. The position of the ultrasound probe and B-Mode settings were adjusted until clear vessel borders with typical M and I lines were visible, and several cine video loops were acquired and saved as video files for off-line analysis. Flow velocity measurements were performed with pulsed wave Doppler. The sample volume was placed at the center of the perfused artery, insonation angle adjusted to <60° and angle correction applied. Image analysis was performed offline with an automated edge detection software (Brachial analyzer, Medical Imaging Applications). The luminal diameter was measured on video loops over several pulsation cycles. Each diameter measurement represents the average diameter over 0.5-1.0 cm vessel area. The maximal diameter was designated systolic diameter and lowest diameter as diastolic diameter. Volumetric flow estimate was calculated as π(mean diameter/2)^2^* mean velocity. Wall shear stress was estimated with 8*μ*mean flow velocity/mean diameter. The viscosity of the perfusion medium (μ) was estimated at 0.035 dyn*s/cm^2^ (10). Intimal media thickness was also measured with this software over the same vessel segment on the far side as the distance between lumen-intima (I line) and media-adventitia interface.

For the functional vasoreactivity assay, syringes containing 1 ml of nitroglycerin solutions were prepared, with concentration ranging from 10^-9^ M to 1 M. These concentrations were calculated so that added to 100 ml of medium contained in the bioreactor, they yielded final concentrations ranging from 10^-11^ and 10^-2^. Nitroglycerin boluses were injected in order of increasing concentration and the reaction of the blood vessel was recorded with a delay of 1 minutes, once the vessel normalized the next bolus was injected.

### Histological preparations

Portions of freshly isolated (control) and cultured arteries were fixed in 4% PFA (Paraformaldehyde, Santa Cruz Biotechnology) overnight (O/N) at 4°C. Fixed samples were either used for whole tissue *en-face staining* or were further processed for histological analysis.

*En-face* staining was performed on samples washed with PBS and permeabilized with 1v/v% TritonX-100 in PBS for 15 minutes. Tissues were blocked overnight at 4°C in 20v/v% goat serum in PBS and then incubated with primary antibody solution (CD31, Abcam-ab28364, 1:50) overnight followed by goat anti-Rabbit Alexa Fluor 488® (Thermo Fisher Scientific) secondary antibody diluted 1:200 and Phalloidin-iFluor 594 (Abcam-ab176757) for 1 hour at 37°C. Nuclei were stained with DAPI (4’,6-diamidino-2-phenylindole, Merck) for 10 minutes at room temperature. Tissues were laid flat and mounted with Fluoromount G (Invitrogen eBioscience Fluoromount G, Thermo Fisher Scientific). Imaging was performed with Nikon Eclipse Ti A1-A confocal laser scanning microscope, obtaining stack images at 40X along the Z-axis.

For histology, tissues were washed with PBS and incubated overnight in 30w/v% sucrose (Sigma-Aldrich) solution in PBS. Following sucrose incubation, samples were embedded in OCT Compound (Agar scientific) in an Iso-Pentane bath cooled using liquid nitrogen. Frozen samples were stored at - 80°C until later use. Samples were cryo-sectioned along the transverse plane, obtaining 5 μm thick sections. Sections were collected on microscope slides in quadruplicate, where sections were separated by at least 250 μm.

For immunofluorescent staining on tissue sections, antigen retrieval was performed in a water bath at 80°C for 30 min in tris-EDTA buffer (10 mM Tris Base, 1 mM EDTA Solution, pH 9.0) followed by blocking for 1 hour at room temperature with 20v/v% goat serum (Sigma-Aldrich) in PBS. Primary antibodies against CD31 (Abcam-ab28364) diluted 1:200, and Proliferating cell nuclear antigen (PCNA, Sigma Aldrich-MABE288) diluted 1: 100 were incubated overnight at 4°C.

Appropriate Goat anti-Mouse and Goat anti-Rabbit Alexa Fluor® (Thermo Fisher Scientific) secondary antibodies 488 and 567 diluted 1:200 were incubated for 1 hour at 37°C. Following secondary antibodies, Human α-Smooth Muscle Actin (SMA) Alexa Fluor® 647-conjugated antibody (R&D Systems-IC1420R) diluted 1:200 was introduced for 1 hour at 37°C. Nuclei were stained with DAPI (Merck) for 10 minutes at room temperature. Incubation with 0.1 w/v % Sudan Black (Sudan Black B, Santa Crus Biotechnology) in 70% ethanol for 10 minutes at room temperature was performed to reduce tissue autofluorescence. Slides were then mounted in Fluoromount G (Invitrogen eBioscience Fluoromount G, Thermo Fisher Scientific) and imaged with Nikon Eclipse Ti A1-A confocal laser scanning microscope. Tile images were obtained at 20X to capture the whole tissue section where possible.

Hematoxylin and Eosin (H&E), and Masson’s trichrome staining (MT) were outsourced to Veterinary School Pathology Centre at the University of Surrey, and performed by an automated staining system (VIP® 6 Vacuum Infiltration Processor). Immunohistochemical assays for Caspase 3 (1:1000, R&D systems, AF835), PDGFR-β (Santa Cruz Biotechnology, sc-374573, 1:100) and Vimentin (1:5000 Dako, Ref M0725, Clone V9, 1:5000) were similarly outsourced to Veterinary Diagnostic Services, School of Veterinary Medicine, University of Glasgow. Primary antibodies were followed by HRP-conjugated secondary antibody and DAB (3,3’-Diaminobenzidine) staining and counter-staining with hematoxylin. Staining was performed in the Dako Autostainer system. Slices were dehydrated and mounted with Cellpath Mounting Media (SEA-1604-00A).

Whole slice imaging was performed on the NanoZoomer 2.0-HT slide scanner by Hamamatsu.

### Image analysis

All acquired images were processed in ImageJ v1.53c (FiJi). Custom macros were composed to facilitate image analysis. Where necessary, vessel wall areas (lumen, intima, media, adventitia) were defined manually and stored as Regions of Interest (ROIs). Resulting ROIs were used to define tissue dimensions and to measure signals specific to the predefined areas.

### Quantification of Cell Activation

The quantification of PCNA was performed on the fluorescent images collected by confocal microscopy. Tile images covering the cross section of the tissue were obtained to measure distribution of PCNA across the tissue for each section (3 sections/sample). PCNA+ nuclei counts were normalized against the total number of DAPI+ nuclei within each area.

To define the spatial distribution of PCNA+ cells, images were converted into distance maps where orthogonal distance of each nucleus from the lumen was calculated. Each distance was further normalized against the thickness of the tunica media, to account for differences in tissue morphology between specimens. Individual distance values were organized into a histogram representing the relative frequency of PCNA+ cells against the relative distance from the lumen.

### Signal quantification in histological samples

All histological samples (IF/IHC) were analyzed the expression of individual markers by averaging the signal across the whole sample. The area presenting the signal was measured and normalized towards the total area.

### Tissue coherence analysis in using Masson’s staining

Histological samples prepared with Masson’s trichomic staining were used to analyze the coherence of the tissue. Three random, uniformly sized, representative regions were selected to perform the analysis. In each region, the area occupied by the tissue versus the empty spaces were measured and normalized towards the total area of measurement.

### Endothelial Coverage quantification

The quantification of endothelial coverage was performed on the fluorescent images obtained by confocal microscopy. To quantify the coverage, the length of CD31-expressing lumen over the total length of the lumen was calculated in vessel cross-sections.

### Fiber alignment

Fiber orientation was quantified in the *en-face* confocal images compressed into a Z-Max projection. In each picture, three random areas were selected and a vector map of the signal was generated using OrientationJ (ImageJ/FiJi plugin) to obtain the local orientation of the signal within the selected area. Resulting vector values were organized into a histogram to represent the relative frequency of angular orientations, where zero is the representative value of fibers perpendicular to the flow direction.

### Statistics

Experiments were repeated in 3 to 5 biological replicates. In some analyses, individual samples have been excluded due to poor quality of the preparation or staining.

Difference among groups were evaluated using one-way ANOVA or Kruskal–Wallis test, based on results from normality tests, followed by Fisher’s LSD post-hoc test (GraphPad Prism 8.1.2). A value of p<0.05 was considered as statistically significant. Data are presented as mean±SD.

## Results

### Design and specifications of the EasyFlow bioreactor insert

The resolution of MultiJet fusion 3D printing enabled the efficient and cost-effective production of the EasyFlow insert, incorporating detailed features, and producing a non-porous surface. The resulting product is biocompatible and autoclavable multiple times. The insert is designed to fit as the lid of a 50 ml centrifuge tube, creating a versatile and self-contained bioreactor for vascular culture (**Figure 1A**). The EasyFlow insert incorporates the following functional features: de-bubbler to remove any air bubbles trapped in the system, 6 inlet/outlet ports creating the potential for 2 separate circulations (inner circulation-going through the vessel sample, and outer circulation-to exchange medium in the tube), and extra access ports for sensors, gas/pressure exchange, and endovascular probes or catheters (**Figure 1A**). The blood vessel is accommodated in an adjustable adaptor, through barb connectors of interchangeable size (**Figure 1A**). Once the bioreactor containing the sample is sealed, it can be connected to an equivalent bioreactor functioning as a medium reservoir and to a peristaltic pump, though medical grade tubing (**Figure 1B and C**). EasyFlow is amenable to Doppler imaging, the probe can be applied directly to the to the exterior wall of the centrifuge tube, enabling longitudinal monitoring of the changes in hemodynamic forces and the structural/functional response of the blood vessel (**Figure 1B**). Additional features are added to the system to improve the performance, such us one-way valves to ensure unidirectional flow, pressure dampeners to regulate the system pressure, and gas exchange filters (**Figure 1C**). Thanks to these features, we ensure that the blood vessel cultured within EasyFlow would experience physiological ‘heart beat’ (56 bpm) and pressure (60-120 mmHg). The media used in our culture was enriched with dextran, to obtain a media viscosity similar to that of human blood (0.035 dyn/sec*cm^2^ or 3.5 cP)

### EasyFlow provides controlled physiological culture conditions

Using values extrapolated by the Doppler/Duplex analysis, we ensured that the carotid arteries within *EasyFlow* were exposed to mechanical parameters closely resembling the physiological arterial hemodynamics. The average diameter of arteries after 7 days of culture under pulsatile flow was 1.24±0.32 mm (‘systolic’) and 1.19±0.30 mm (‘diastolic’). The ‘peak systolic velocity’ was 52±13 cm/s and mean velocity was 44±8 cm/s, resulting in an estimated volumetric blood flow of 33±16 ml/min (11) and wall shear stress of 10.1±2.8 dyn/cm^2^ (12)) (**Table 1**). The average IMT was 0.32±0.08 mm. Overall repeated measurements of vessel diameter showed that there was an average deviation (AD) of 0.00±0.03 mm between 2 independent automated measurements of 4 vessels. See **Table 1** for repeatability of other parameters. In summary, these characteristics support that *EasyFlow* provides reproducible *ex vivo* hydrodynamic conditions that resemble physiological hemodynamics *in vivo*.

**Table 1.** Characteristics of culture within the EasyFlow and their repeatability. Mean value (MVal) and average deviation (AD) of vessel parameters in flow culture at day 7. IMT= intima media thickness, PSV = peak systolic velocity, MV= mean velocity, WSS = wall shear stress, Systolic= maximal pulsatile diameter, Diastolic= minimal pulsatile diameter (n=4). Bland Altman plot shows the consistency of repeated measures within the samples.

| | MVal±SD | AD±SD |
| --- | --- | --- |
| Systolic diameter (mm) | 1.24±0.32 | 0.00±0.02 |
| Diastolic diameter (mm) | 1.19±0.30 | 0.00±0.04 |
| IMT (mm) | 0.32±0.08 | -0.01±0.02 |
| PSV (cm/s) | 52±13 | 2±3 |
| MV (cm/s) | 44±8 | 0±4 |
| Volumetric flow (ml/min) | 33±17 | 0±2 |
| WSS (dyne/cm<sup>2</sup>) | 10.1±2.8 | 0.0±1.3 |

### EasyFlow culture preserves extracellular matrix organization and tissue homeostasis

Our finding confirmed that EasyFlow provides physiological culture conditions to arterial tissues. Next, we tested how the EasyFlow environment impacted on cellular physiology and ultrastructural components, by performing *en-face* staining, immunofluorescence imaging and histology in porcine carotid arteries segments cultured in EasyFlow or in static culture for 7 days, and compared them to their freshly isolated controls.

Hematoxylin and eosin staining revealed that the overall structure of the tissue was preserved during culture under flow conditions. In particular, the vessel wall layers appear unaltered, and no remodeling was observed (**Figure 2A**). Samples cultured in static presented a distinct displacement of the intima, and loss of structure in the media and adventitia (**Figure 2A**).

**Figure 2.**
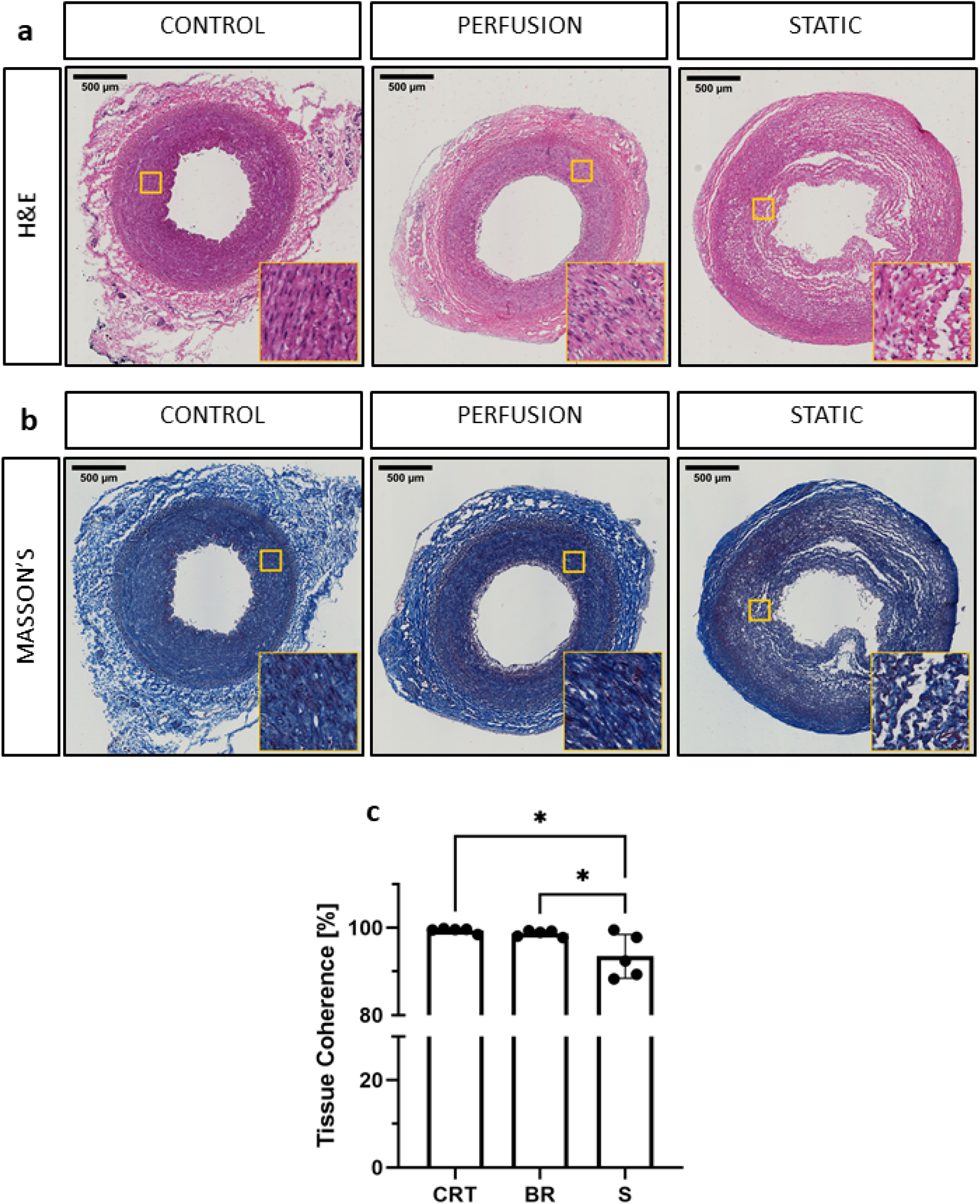
Histological evaluation of porcine carotid arteries cultured in EasyFlow or static conditions. Hematoxylin and Eosin staining (H&E, a) and Masson’s Trichomic staining (MASSON’S, b) of freshly isolated carotid artery tissue (CONTROL) and samples cultured for 7 days in EasyFlow (PERFUSION) or static culture (STATIC). Scale bars= 500 μm. Quantification of tissue coherence is displayed in the bar graph (c), n=5. All data are mean±SD. One-Way ANOVA analysis was performed, *=p ≤ 0.05.

Extracellular matrix provides the mechanical support and elasticity necessary for blood vessels to perform their function, cell apoptosis and abnormal activation may affect this structure, weakening the vessel wall. We visualized the collagen content by Masson’s trichrome staining and confirmed the maintenance of the extracellular matrix organization in tissues cultured in EasyFlow, while static culture produced a visible loss of tissue integrity (**Figure 2B**). These observations were reflected in the quantification of tissue coherence which indicated an increase in tissue disarrangement in static cultured samples (**Figure 2C**).

Immunofluorescence staining revealed the finer changes in cell distribution. Both endothelial coverage of the lumen and expression of smooth muscle actin in the media were slightly reduced in the static condition, as compared to the flow culture, although not to a significant level (**Figure 3A-C**). These modest changes were reflected by the histological analysis of apoptosis and proliferation. Staining for the apoptosis marker Caspase-3, indicated no significant increase in cell death following culture, although a trend is clearly visible in 3/5 static culture samples (**Figure 3D**). Quantification of the overall percentage of PCNA+ cells revealed a slight increase in proliferation over culture, but no significant differences compared to the control (**Figure 3E**). We also detected a decrease in Vimentin+ cells in the static culture, potentially reflecting a loss of adventitia (**Figure 3F**), and an overall maintenance of PDGFRβ+ expression (**Figure 3G**).

**Figure 3.**
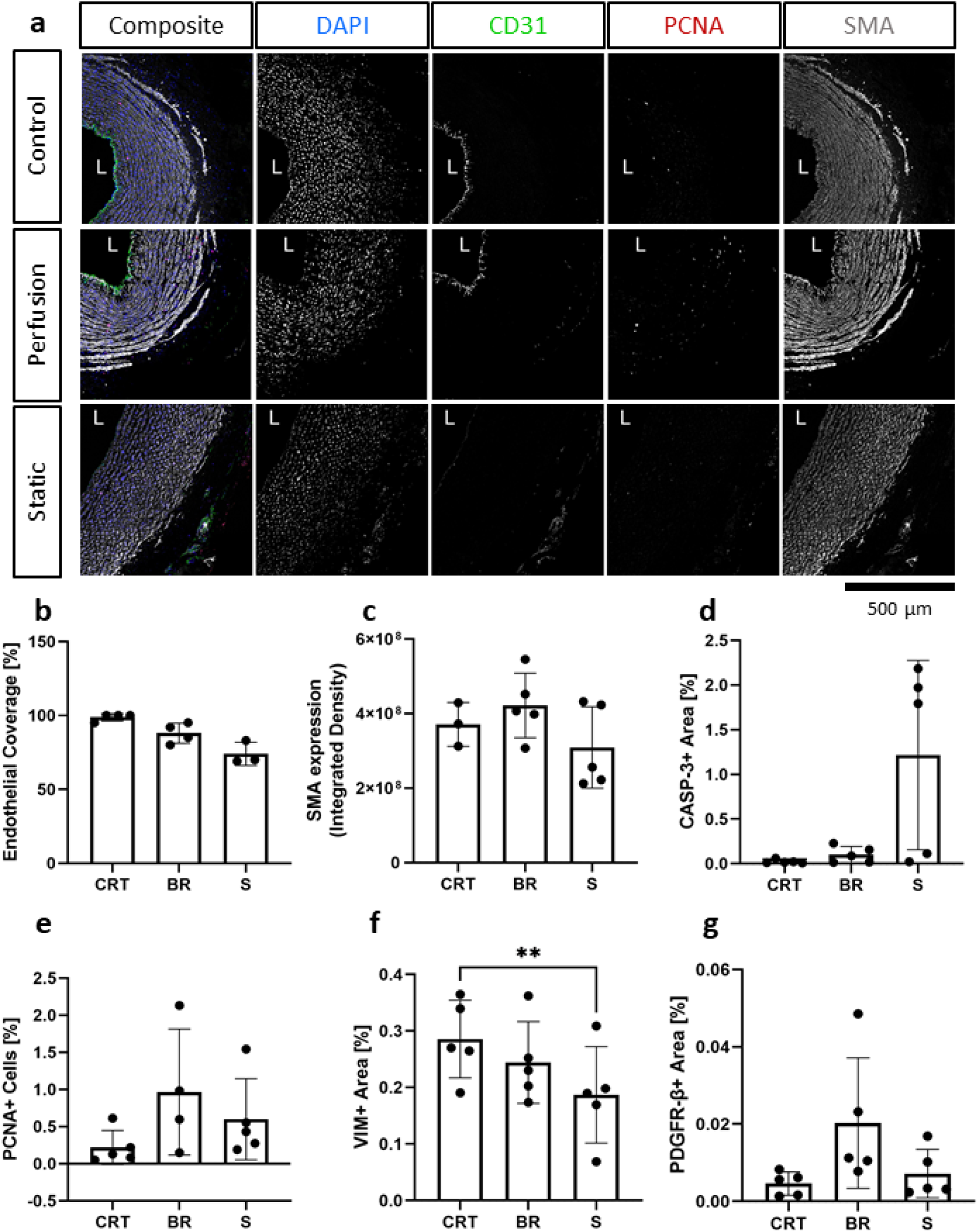
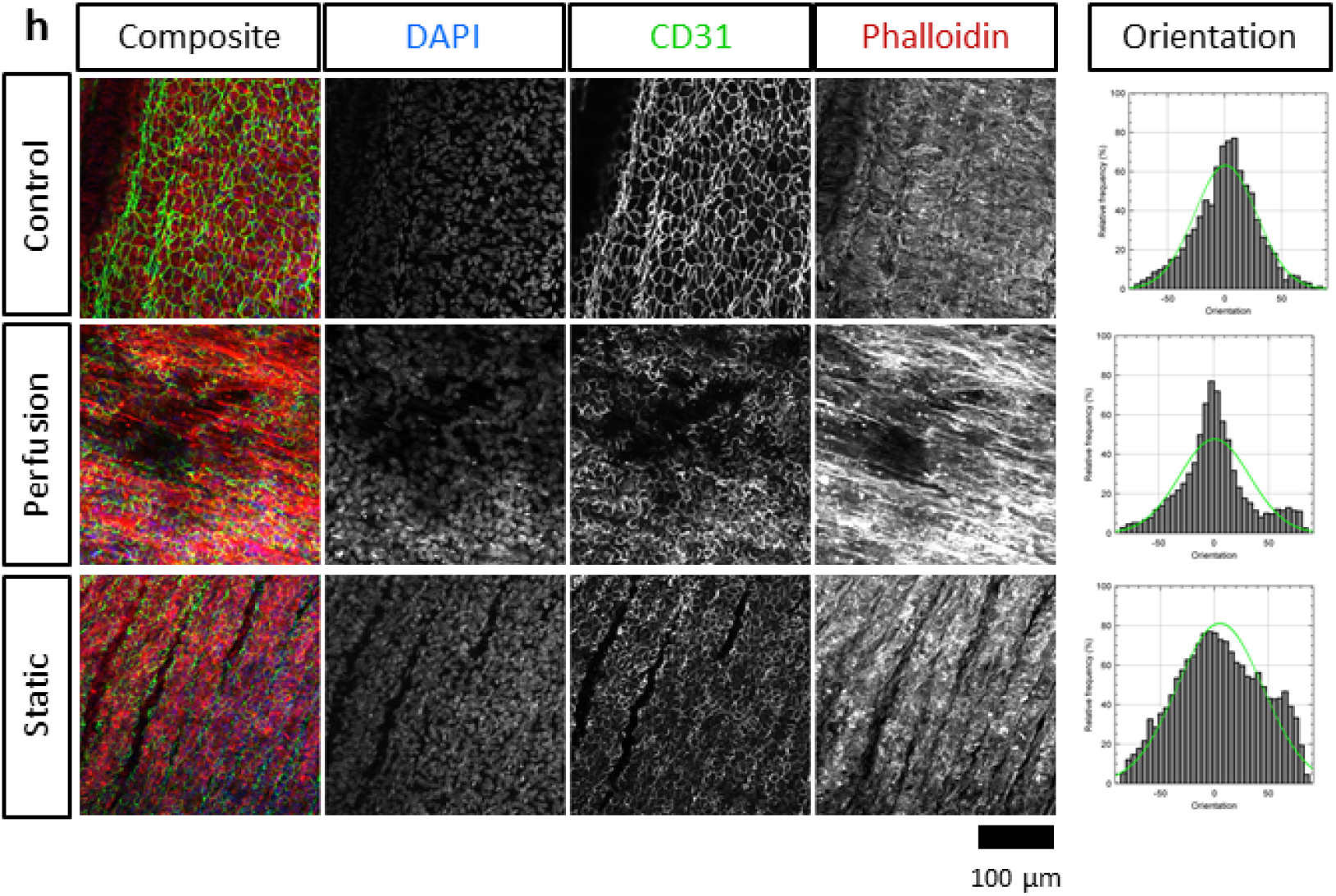
Immunofluorescence analysis of cultured carotid arteries indicates better tissue preservation in flow. Immunofluorescence staining of freshly isolated arteries (Control/CRT), EasyFlow (Perfusion/BR) and static cultured arteries (Static/S) at day 7, showing expression of endothelial marker CD31, smooth muscle marker smooth muscle actin (SMA), and proliferation marker proliferating cell nuclear antigen (PCNA). Nuclei are stained with DAPI. Representative confocal images are displayed in composites and by individual marker in montage (a). Lumen is identified (L), Scalebar= 500 μm. Quantifications of lumen coverage (b), SMA intensity (c), apoptosis (caspase 3, casp-3, d), proliferation (PCNA, e), vimentin (VIM, f) and platelet-derived growth factor receptor beta (PDGFRβ, g) are displayed in the bar graphs, n=3-5. All data are mean±SD. One-Way ANOVA analysis was performed, *=p ≤ 0.05, **=p≤0.01. *En-Face* representative images showing endothelial coverage (CD31), stress fiber (Phalloidin), and nuclei (DAPI) in samples cultured for 7 days in perfusion or static after injury (h). Fiber alignment is quantified as vector distribution and plotted in histogram representing the relative frequency (y) against the angle (x), where zero is perpendicular to the flow direction (Orientation). An ideal normal distribution of values is represented in green over the histogram (h). Scalebar= 100 μm.

Next, *en-face* staining was performed to better visualize endothelial coverage and media organization. Overall, *en-face* staining revealed a good luminal coverage in both culture conditions (**Figure 3H**), however culture in flow produced more organized structures in the media, which were comparable to the native tissues, while static culture conditions led to fiber disarray (**Figure 3H**). Taken together these results indicate that culture of porcine arteries in physiological flow conditions helps maintaining appropriate cellular organization and structural integrity.

### EasyFlow cultures are amenable to endovascular balloon procedures

The study of vascular pathology *ex vivo* has the potential to reduce and replace the number of animals used as models, providing results that are more physiologically relevant and may have scope to improve the patient care.

We performed balloon injury in porcine carotid arteries before culturing them for 7 days in flow conditions. An important advantage of the EasyFlow system is the ability to monitor the structural changes and functionality of the vessel longitudinally, by Doppler ultrasound. Here, we used it to determine the luminal diameter and intima media thickness and blood flow within the artery, providing a closer estimate of the hemodynamic environment within the vessel. We also investigated functional response of the vessel wall to increasing doses of the endothelium-independent vasodilator nitroglycerin. Ultrasound showed that at day 3 the diameter of the vessels was significantly greater after angioplasty as compared to control (systolic: 1.17±0.52 mm vs 2.79±0.12 mm, p=0.026, diastolic: 1.09±0.52 mm vs 2.47±0.04 mm, p=0.044). While control retained the diameter at day 7, injured arteries significantly decreased diastolic diameter (control:1.19±0.30 mm, p=0.779; angioplasty: 1.44±0.60 mm, p=0.041). There were no significant changes in blood flow (**Figure 4A and B**). We leveraged on our ability to monitor the blood vessel in culture by ultrasound to quantify tissue remodeling using clinically relevant readouts. Intima-Media thickness (IMT) is considered a gold standard for the prediction of subclinical vascular remodeling and propensity to atherosclerosis in patients. We performed a preliminary analysis by measuring IMT in B-mode images, and showed that after injury there was an increase in IMT. While IMT remained stable between day 3 and 7 in control (0.27±0.04 mm and 0.32±0.08 mm, p=0.355), it was decreased as compared to control in angioplastied arteries at day 3 (0.15±0.01) and increased at day 7 (0.38±0.06 mm) (**Figure 4A and B**). We also compared the vascular response of the injured and control arteries cultured in flow. This identified a loss of 60-70% endothelium-independent vasodilator response upon stimulation with increasing doses of nitroglycerin (**Figure 4C**).

**Figure 4.**
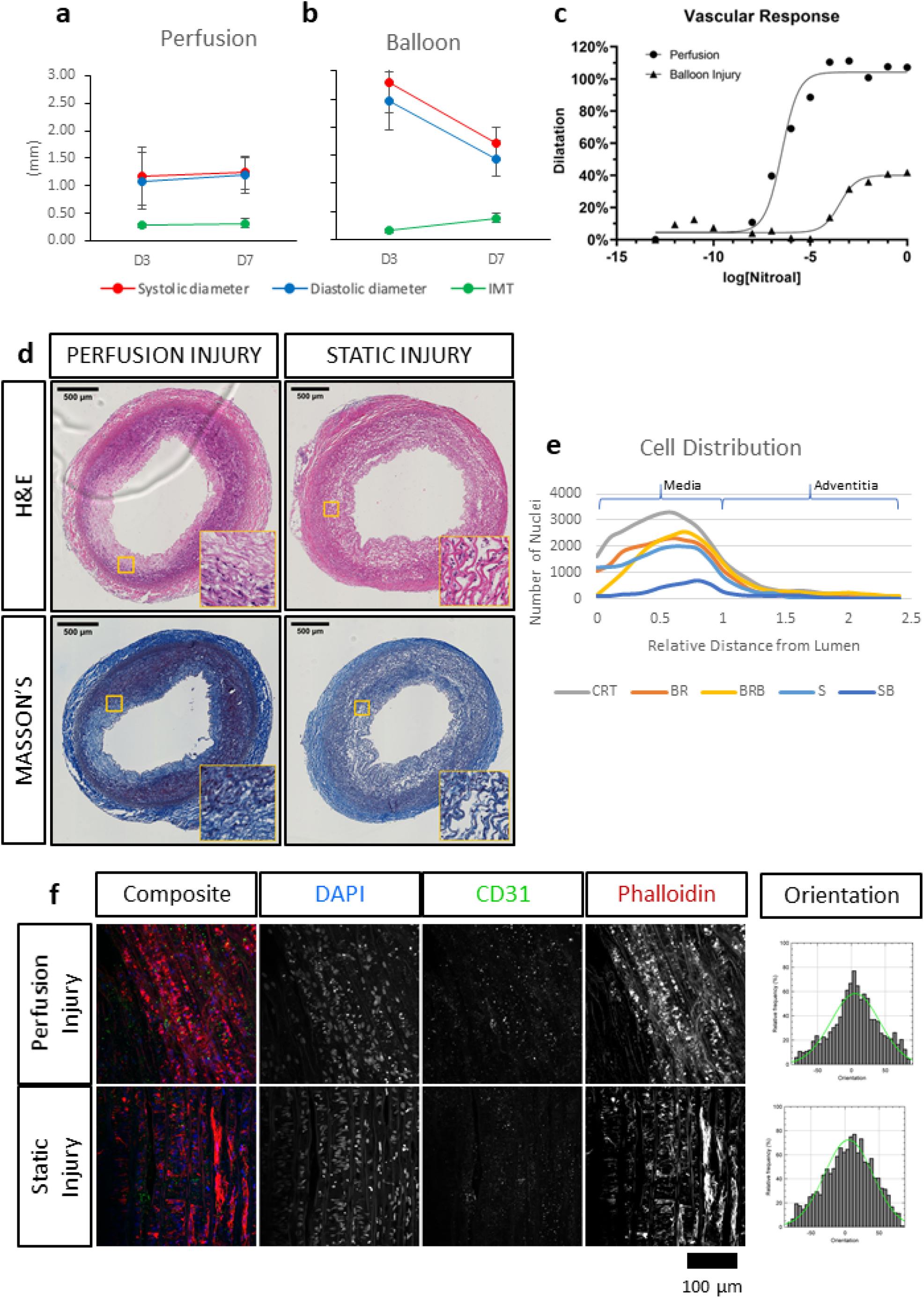
Ultrasound and immuno-histological evaluation of angioplastied carotid arteries after culture reveal functional and structural impairment. Healthy (Perfusion, a) and angioplastied (Balloon, b) arteries were cultured for 3 and 7 days in EasyFlow and their systolic and diastolic diameter, the intima-media thickness (IMT), and their dose-response to endothelial-independent vasodilator nitroglycerin (c) were quantified by Doppler ultrasound, n=2-4. Injured vessels were cultured in EasyFlow (Perfusion Injury) or in static conditions (Static Injury) for 7 days and analyzed by Hematoxylin and Eosin staining (H&E) and Masson’s Trichomic staining (MASSON’S) (d). Scale bars= 500 μm. Distribution of DAPI+ nuclei across the vessel wall is quantified in freshly isolated arteries (CRT), healthy and injured arteries cultured in EasyFlow (BR and BRB, respectively) or static conditions (S and SB, respectively) (e). *En-Face* representative images showing endothelial coverage (CD31), stress fiber (Phalloidin), and nuclei (DAPI) in samples cultured for 7 days in perfusion or static after injury (f). Fiber alignment is quantified as vector distribution and plotted in histogram representing the relative frequency (y) against the angle (x), where zero is perpendicular to the flow direction (Orientation). An ideal normal distribution of values is represented in green over the histogram (f). Scalebar= 100 μm.

Next, we compared the ultrastructural effect of culturing injured arteries in static and flow conditions by histology, immunofluorescence and *en face* staining. As expected, balloon injury determined a significant loss of cells and disorganization in the extracellular matrix in the intima and the medial layers, as shown in the H&E and Masson’s trichrome staining (**Figure 4D**). The effect of the injury was dramatic in tissues cultured in static, likely due to the poor preservation in these conditions. In arteries cultured in flow conditions, the damaged media after injury corresponded to 21.7±11.6% of the total vessel wall. In order to visualize the spatial effect of the injury, we quantified the distribution of nuclei within the vascular wall in culture, with or without injury. Results show that in non-injured samples, the static culture determined a baseline reduction of cell number in the media, and that the balloon injury almost completely abolished cells in these samples (**Figure 4E**). In flow conditions, we observed a striking loss of cells close to the lumen upon injury, and an accumulation of cells in the distal part of the media, at the junction with the adventitia (**Figure 4E**). *En face* staining confirmed that balloon injury provoked extensive endothelial denudation in both culture system and that the alignment of the F-actin fibers was lost in the flow culture, as a result of extension damage (**Figure 4F**). Immunofluorescence analysis confirmed the significant loss of endothelial coverage, and a mild reduction in SMA expression in vessels cultured in flow after injury, which is likely due to large areas of damage produced in the media (**Figure 5A-C**).

**Figure 5.**
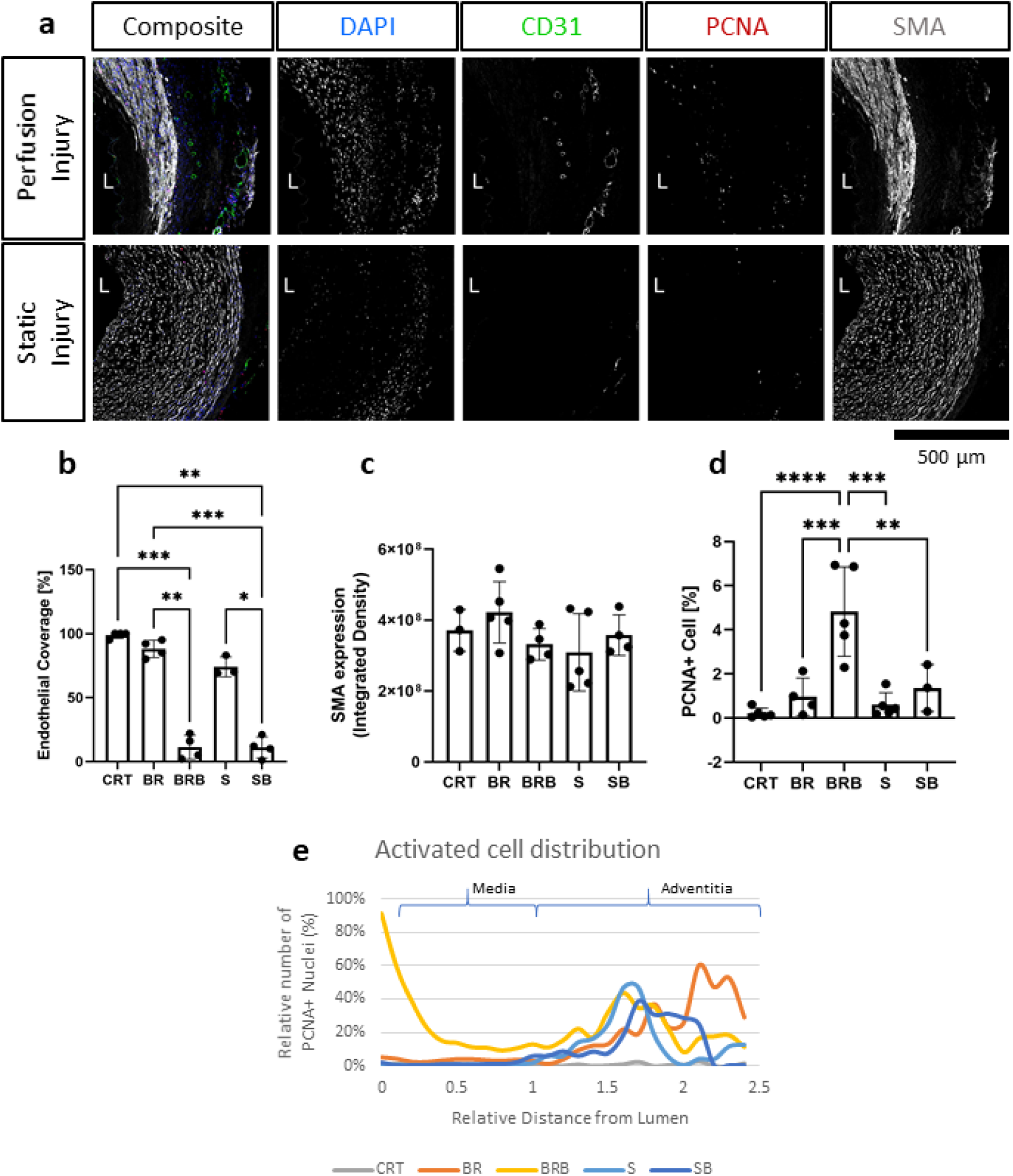
Immunofluorescence evaluation reveals tissue injury and activation following balloon angioplasty and culture in flow. Following injury, tissues were cultured for 7 days in either perfusion (Perfusion Injury) of static conditions (Static Injury) and then stained for the endothelial marker (CD31), smooth muscle marker (smooth muscle actin, SMA), and proliferation marker (proliferating cell nuclear antigen, PCNA). Nuclei are stained with DAPI. Representative confocal images are displayed in composites and by individual marker in montage (a). Lumen is identified (L), Scalebar= 500 μm. Quantifications of lumen coverage (b), SMA intensity (c), and proliferation (PCNA, d) are displayed in the bar graphs, n=3-5. All data are mean±SD. One-Way ANOVA analysis was performed, *=p≤0.05, **=p≤0.01, ***=p≤0.001, ****=p≤0.0001. Distribution of proliferating cells (PCNA+) in relation to their relative distance to the lumen is plotted for freshly isolated arteries (CRT), healthy and injured arteries cultured in EasyFlow (BR and BRB, respectively) or static conditions (S and SB, respectively) (e).

Taken together these results show that EasyFlow cultured arteries subjected to endovascular damage present areas of cell loss and functional impairment, but maintain overall structural integrity.

### Injured arteries cultured in EasyFlow present signs of remodeling

*In vivo*, vascular injury leads to extensive apoptosis and cell replacement, which triggers the activation of the vascular wall progenitor niche and the synthetic switch of vascular smooth muscle cells. Pathological remodeling generally follows, with the formation of fibrosis, neointimal growth or atherosclerotic plaques.

We analyzed the arteries cultured for 7 days in static or EasyFlow post-injury. Overall proliferation was uniquely increased in the injured samples cultured in flow (**Figure 5D**), and the PCNA+ cells were mainly localized at the interface between the media and the intima, and in the luminal side of the media (**Figure 5A and E**). Proliferating cells mainly co-expressed SMA, and were found largely in undamaged area of the media (**Figure 6Ai**), however we also detected a unique population of proliferative SMA+ cells, separated from the injured media, and located beneath the lamina elastica (**Figure 6Aii**). It was also possible to identify the occasional proliferative CD31+ cell in the intima (**Figure 6Ai**) and the adventitia (**Figure 6Aiii**). Organized microvascular structures can also be observed penetrating the media after injury (**Figure 6A**, red arrowhead), as well as infiltrating mononucleated cells which could potentially represent resident immune cells (**Figure 6A**, orange arrowhead).

**Figure 6.**
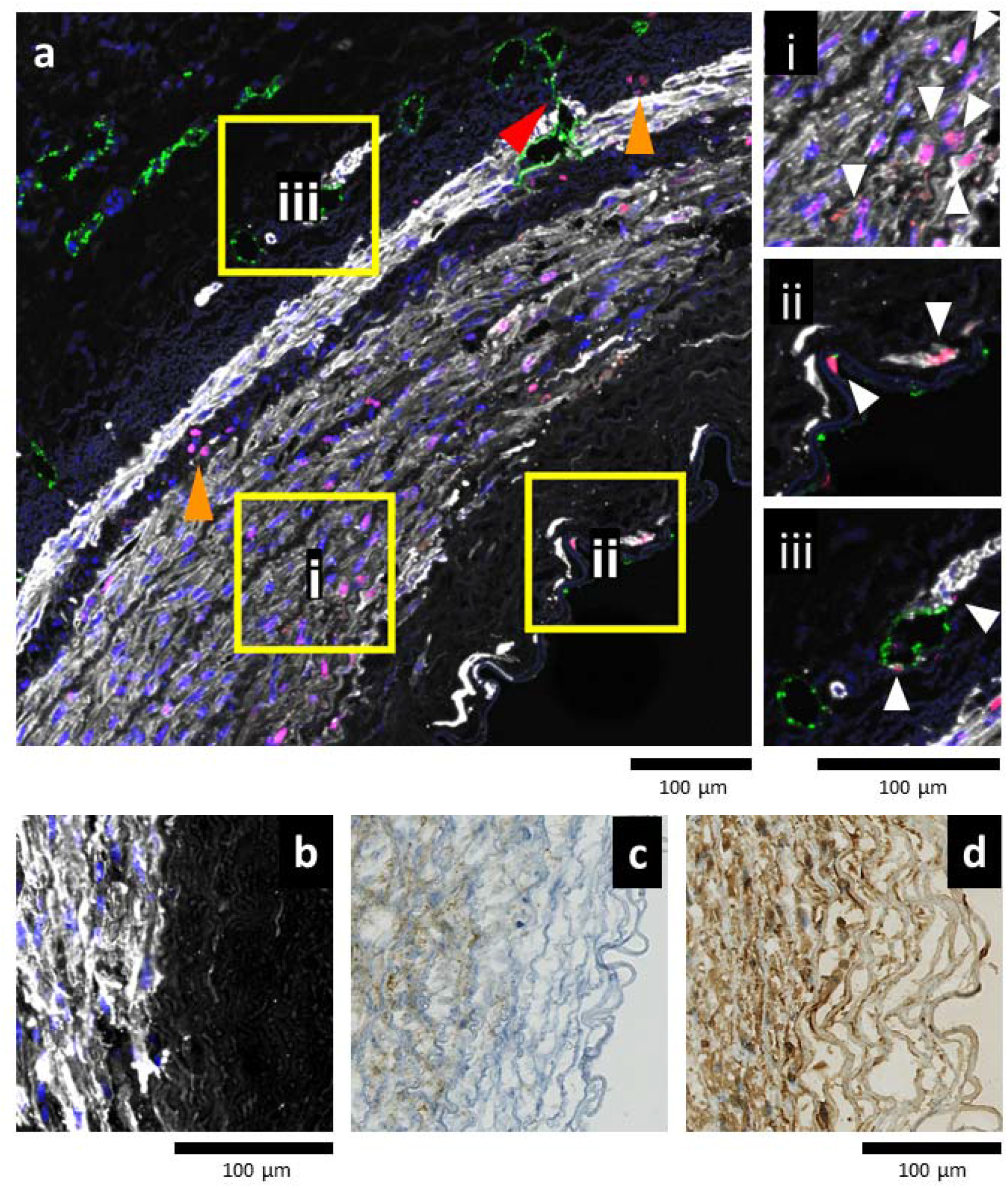
Balloon angioplasty stimulates early remodeling in EasyFlow cultured arteries. Representative confocal images of the arterial wall following injury and flow culture (a) reveals proliferation (proliferating cell nuclear antigen, PCNA, magenta) of smooth muscle cells (smooth muscle actin, SMA, white) in the proximity of the injured media (ai). In the lumen area, individual proliferating (PCNA+) smooth muscle cells (SMA+) and endothelial cells (CD31+, green) are located either side of the lamina elastica (aii). In the adventitia, proliferative endothelial cells (CD31+) line the microvasculature of the vasa vasorum (aiii). White arrowheads indicate proliferative cells. Across the vessel wall, groups of infiltrating cells display PCNA+ nuclei (orange arrowheads), and small blood vessels sprout from the adventitia towards the media (red arrowhead). Nuclei are stained with DAPI, Scale bar= 100 μ m. Smooth muscle cells near the injury area display high expression levels of SMA (white, b), Vimentin (c) and PDGFRβ (d). Scale bars= 100 μ m.

Immunostaining also revealed that in the border zone near the medial damage, smooth muscle cells strongly expressed SMA, Vimentin and PDGFRβ (**Figure 6B-D**). This indicates that smooth muscle cells in these areas were provoked into the synthetic phase by the injury.

Our results showed that culture under flow better recapitulate the early characteristics of post-injury vascular remodeling, as compared to static culture, and that early hallmarks of activation and remodeling can be identified in flow cultures by histology and ultrasound imaging.

## Discussion

In this study we presented a 3D printed bioreactor that is economical to produce, easily transferable to other labs and enables small volume cultures and multiplexing. EasyFlow cultures can be imaged longitudinally by Doppler ultrasound, enabling the monitoring of flow conditions and tissue integrity in a non-invasive manner. Using EasyFlow bioreactor, porcine carotid arteries can be cultured in physiological conditions closely resembling the human arterial circulation. When compared to static cultures, tissue integrity and organization was superior in flow cultures. Endovascular balloon injury provoked intimal denudation and media damage, also detected by Doppler ultrasound imaging. After injury, samples cultured in EasyFlow showed increased proliferation in the media and adventitia, and overexpression of smooth muscle markers near the injury. A small population of highly proliferative SMA+ cells was also observed at the intima/media interface. Samples cultured in static were instead significantly compromised and did not show signs of activation.

The study of vascular physiology and pathology is often pursued with the use of moderate severity animal models such as carotid ligation or wire injury, in some case presenting genetic mutations to emulate specific co-morbidities such as diabetes (db/db) and hypercholesterolemia (ApoE-/-) (3). These *in vivo* experiments are routinely supported by *in vitro* work to help elucidate mechanistic insights, using static monocultures of vascular cells, which struggle to recapitulate the physiologically relevant mechanical cues and the interaction with other cell types.

Despite these limitations, these models delivered great insights in the understanding of the mechanisms of vascular homeostasis and the patterns associated with common diseases, such as atherosclerosis and restenosis (13). The striking physiological differences have however determined disappointing outcomes for some of the therapeutic approaches in clinical settings, sparking new interest in large animal models for these diseases (14). However, ethical and economic implications associated with this type of animal work demands new solutions.

Vascular bioreactors have been developed both at commercial level and in-house to provide a suitable physiological environment to maintain blood vessels in culture or enhance the maturation of tissue engineered constructs.

In this study, we present a 3D printed bioreactor that enables the culture of blood vessels within a 50 ml centrifuge tube, enabling small volume cultures and multiplexing. EasyFlow’s open-access design allows simple uptake by other labs, and the adjustable vessel compartment enables customization to different vessel sizes and tissue sources by simply exchanging the barbed connectors and adjusting the length adapters.

Dynamic bioreactors supply culture conditions that are as close as possible to the *in vivo* environment experienced by the blood vessel in its natural location. Common carotid arteries experience a mean flow velocity of 15-43 cm/s (end diastolic-peak systolic) and a shear stress in the range of 6-21 dyne/cm^2^ (end diastolic-peak systolic) (15,16). EasyFlow culture conditions were calculated to provide a physiological environment in the culture, and encompassed a mean flow velocity of 44±8 ml/min, and a mean wall shear stress of 10.1±2.8 dyne/cm^2^, which are comparable to the arterial flow forces *in vivo*. Pump revolution rate was adjusted to 56 rpm to simulate resting heart rate and continue pressure monitoring confirmed a physiological range of 60-120 mmHg, which is compatible to the average healthy blood pressure in the human population. The resulting volumetric flow imposed is within the range measured in equivalent vessels *in vivo* (11). In addition, we modified the medium composition to simulate the viscosity of blood (3.5-5.5 cP), this parameter closely affects the shear stress experienced by vascular cells in flow, according to the Hagen-Poiseuille equation (16). Our in-depth consideration and characterization of the conditions and forces acting on the vessel in culture is uncommon in previously published studies (9,17-19). Importantly, these forces were initially estimated by calculation, but also verified within the cultured blood vessel by Doppler ultrasound imaging. While few examples exist in literature, we present a uniquely exhaustive application of the Doppler ultrasound method to a bioreactor culture. This analysis enables direct comparison with patient studies, and allows to visualize vessel structure and functionality over time (20).

When compared to static culture, EasyFlow cultured arteries demonstrated an overall better maintenance of tissue integrity and structure. In particular, the organization of the smooth muscle cells and the extracellular matrix were closer to the native tissue. Interestingly, we did not detect a critical difference in lumen coverage, apoptosis or activation of the vessel wall in the two culture systems. This is not completely unexpected as the static culture was designed to replicate the vessel ring culture, with tissue specimens of 2-5mm in length (5,21-23). Previous studies have demonstrated that static culture of rings can be extended up to 56 days, and can be induced to form neointima post-injury, although we did not detect such a growth in our samples (22). Samples cultured in flow were instead much longer (2-4 cm), and would not likely have survived in static culture due to limited oxygen and nutrient diffusion.

After verifying the feasibility of a physiological culture using EasyFlow, we assessed the effect of injury in the flow cultures. We delivered the injury before setting up the static or flow culture, but it is important to note that, thanks to the multiple access ports available on EasyFlow, the catheter-mediated balloon injury could potentially be implemented on the mounted sample at any point of the culture, without disturbing the system.

*In vivo*, mechanical damage and flow disruption, accompanied by damage to the medial layer (for example by large balloons), results in smooth muscle cells activation, proliferation and migration into the intima layer, contributing to the thickening of the vessel wall and effectively narrowing the lumen (restenosis) (24,25). Adventitial cells, such as resident progenitor cells and pericytes, macrophages and fibroblasts also become activated after injury, releasing cytokines and growth factors and migrating towards the lumen; their contribution to neointimal lesions and the mechanisms associated with it are still highly debated (26,27).

Following balloon application and flow culture, we observed significant damage to the intima and media layers of the tissue, which is comparable to what reported in similar animal models where intima denudation and early media apoptosis drive the regeneration process (28). In addition, the surviving smooth muscle cells in the media responded to the injury by initiating a robust proliferative response. This is akin to the results obtained after 1 week in rabbit iliac arteries, even in terms of percentage of proliferative cells (roughly 4%) (29). Other hallmarks of early remodeling included the upregulation of contractile proteins in the smooth muscle cells adjacent the injury, and the proliferation of cells in the adventitia.

Interestingly, we did not observe the formation of neointima. This is likely due to the limited timeline of the experiment, as usually neointima forms *in vivo* after at least 2 weeks (29). Previously published work demonstrated that flow culture of porcine arteries is suitable for detection of restenosis 7 days after placement of a stent, this could be due to the smaller damage provoked by the stent as compared to the balloon, and the permanence of the ‘stimulus’ as the stent remains in place after intervention (6). Similarly, low impact stimulation such as non-physiological low flow and intima denudation reportedly led to formation of neointima in vessels cultured under flow (6,9). After balloon injury, the loss of contractility in the vessel wall was quantifiable in the ultrasound profile at baseline levels and following challenge with vasoconstrictors. In addition, we were able to quantify an increase in IMT, a commonly used clinical indicator of remodeling (30). In samples cultured for 7 days after vascular injury IMT was increased, a measure that is associated with increased risk of stenosis-atherosclerosis in patients.

One critical characteristic of EasyFlow is that its open-source 3D printed nature enables easy adoption by other labs, and encourages reproducibility. This is fundamentally different from previously proposed systems, which were either based on commercially available bioreactors or in house-built systems (6–9). These systems are either extremely expensive or almost impossible to replicate, including specialized glassware and limited description of the system. In addition, most commercial and in house systems are large and complex, and this has implications in the volume of reagents and physical space necessary, and therefore impact on the capacity to culture more than one sample at one time.

In our design, we attempted to address these and other limiting factors which are impeding the uptake by other labs and reproducibility of culture: (a) we created an open-source file for 3D printing of EasyFlow and (b) the cost of printing is <£100 per piece, (c) EasyFlow is autoclavable and re-usable. Our design enables (d) low volume culture (<100ml), ideal for pharmacological studies, and (e) multiplexing due to the small size of the system.

This proof of principle study demonstrated its suitability for physiological culture of large animal arteries and the study of pathological processes and highlighted the scientific relevance of our *ex vivo* model and the versatility for both basic and interventional studies.

## Author Contributions

RSM conceived and planned the experiments and analyzed the data. DM and CH provided key experimental expertise. RSM and PC wrote the manuscript and prepared the figures with support of CH. PC, CH and JMV conceived the original idea and supervised the project.

## Acknowledgments

The authors wish to thank the Veterinary Pathology Centre at the University of Surrey’s School of Veterinary Medicine for histology services and the Histology Research Service, Veterinary Diagnostic Services, University of Glasgow. We also thank Drs L Dixton, A Reis and M Henstock from the Pirbright Institute (Pirbright, UK) for their support in procuring the animal tissues, and the Department of Biochemical Sciences at the University of Surrey, especially the technical team, for their continuing support.

## Data Availability Statement

The datasets for this study will be made available on Zenodo upon publication and will be shared upon reasonable request.

